# CRISPRi Screening Identifies SON and MAP4K1 as Regulators of Type III Cytokine Expression in Innate Lymphoid Cells

**DOI:** 10.1101/2025.08.15.670561

**Authors:** Rachel A. Brown, Andrew W. Dangel, Ankita Saini, Patrick L. Collins, Marco Colonna, Eugene M. Oltz

## Abstract

The cytokines interleukin (IL)-22 and IL-17 are secreted by innate and adaptive immune cells to drive “type III” responses that protect against extracellular pathogens, promote mucosal barrier integrity, and foster microbiota homeostasis. However, dysregulation of IL-22 and/or IL-17 contributes to autoimmunity, chronic inflammation, and malignancy. Thus, a deeper understanding of mechanisms regulating type III cytokine production could provide new therapeutic targets for a spectrum of immune-mediated diseases. Toward this goal, we performed a genome-wide CRISPR inhibition (CRISPRi) screen to identify factors that regulate IL-22/IL-17 expression in a murine type III innate lymphoid cell (ILC3) model, MNK3, following stimulation with IL-23 and IL-1b. In addition to previously known regulators of type III cytokines, including IL-23 receptor components IL23R and IL12RB1, the screen identified a large set of new factors that either potentiate or attenuate expression of IL-22 and/or IL-17. A subset of these novel factors was chosen for validation, from which two were selected for further study. The nuclear protein, SON, which binds both DNA and RNA, impaired expression of IL12RB1 at the levels of de novo transcription and RNA processing. The second, MAP4K1 (HPK1), is a serine/threonine kinase that is required for IL-22 but not IL-17 expression. Depletion of MAP4K1 in MNK3 also enhanced expression of the type I cytokine, IFNg, which was co-expressed with IL-17, a phenotype reminiscent of pathogenic Th17 cells. Together, results from the CRISPRi screen broaden our understanding of the factors involved in type III immune responses and offer new targets for modulating IL-22/17 expression.

## INTRODUCTION

Optimal regulation of cytokine expression is crucial for immune cell function. Interleukin (IL)-22 and IL-17A/F are the signature cytokines expressed by type III immune cells. Together, they protect against extracellular pathogens, promote wound healing, and drive epithelial cell growth (1, 2, 3). These cytokines are often produced by cells in response to agonists such as IL-1b, IL-23, IL-6, and aryl hydrocarbon receptor (Ahr) ligands (1). Naïve CD4+ T cells that polarize into the T helper (Th)17/22 subset, as well as their innate counterpart, type 3 innate lymphoid cells (ILC3s), are prominent sources of IL-17/22 (3, 4, 5). ILC3s localize to barrier sites (e.g., mucosal and skin), play key roles in microbiota homeostasis, and are vital to early immune responses (2, 4, 6, 7). IL-1b and IL-23 are capable of inducing type III cytokines independently, but act synergistically, and IL-23 signaling is needed for their sustained expression (8, 9, 10, 11, 12). Factors downstream of IL-1b and/or IL-23 receptors (IL1R1/IL1RAP and IL12RB1/IL23R, respectively), including STAT3, RORyt, MAPK, and NF-kB, cooperate to drive IL-17/22 expression, while negative regulators include transcription factors c-Maf and Aiolos and the RNA binding protein ZFP36/TTP (1, 4, 12, 13, 14, 15).

Despite their beneficial properties, chronic expression of type III cytokines contributes to an array of inflammatory conditions, including psoriasis, rheumatoid arthritis (RA), and inflammatory bowel disease (IBD), all of which are characterized by dysregulation of Th17 cells and ILC3s (4, 5, 6, 16). As such, cytokine signaling pathways have become attractive targets for therapeutics, with biologics and small molecule inhibitors targeting the IL-23/-17/-22 axis showing efficacy in psoriasis, Crohn’s Disease, and other autoimmune conditions (17, 18, 19). Selective inhibition of immune system pathways is important to avoid broad immunosuppression and accompanying increases in the risk of opportunistic infections (20, 21). While the number of cytokine-modulating medications has grown, continued research is needed as not all patients respond to current therapies, and there remain numerous cost and logistical barriers (19, 22, 23). Moreover, potential applications for cytokine-based therapies are expanding, such as for oncology (4).

Although many of the key signaling intermediates and transcription factors required for type III cytokine expression are well-studied, others likely have yet to be discovered. Identifying additional key regulators of IL-22 and Il-17 would enhance the ability to modulate their expression and link mutations with diseases. To discover genes involved in various phenotypes, one powerful method of broad, unbiased testing is the use of genome-wide clustered regularly interspaced short palindromic repeats (CRISPR) screening (24, 25, 26). Next generation CRISPR methods utilize a catalytically dead Cas9 (dCas9) in conjunction with transcriptional inhibitors (CRISPRi) or activators (CRISPRa) to suppress or overexpress target genes, respectively (27, 28, 29). In CRISPRi, the dCas9 is fused with a potent chromatin repressor, often a Kruppel associated box (KRAB) domain, which when guided to promoters by an sgRNA, suppresses gene expression, leaving DNA sequences undisturbed (24).

Because cytokine production is a readily detectable phenotype, the application of CRISPR screens to investigate relevant regulatory circuits is an exciting opportunity for driving therapeutic discoveries. Here, we perform CRISPRi screening in an ILC3 model after stimulation by the classical type III agonists, IL-1b/23, to identify factors that impact expression of IL-22 +/- IL-17. Among other novel and known factors, we discovered that the kinase MAP4K1 and the DNA/RNA binding protein SON regulate type III cytokine expression, impacting distinct pathways. Specifically, we found that SON regulates expression of the IL-23 receptor component, IL12RB1, and that depletion of MAP4K1 attenuates IL-22 but is dispensable for expression of IL-17F/A.

## MATERIALS AND METHODS

### Cells

HEK293T cells were cultured in DMEM (Gibco) supplemented with 10% fetal bovine serum (Sigma), 1X Penicillin-Streptomycin (Gibco), and 50 µM 2-mercaptoethanol (Sigma) in 5% CO2 at 37°C. MNK3 cells (NIH-Swiss genotype) were cultured in DMEM supplemented with 10% fetal bovine serum, 1X Penicillin-Streptomycin, 10 ng/ml mouse recombinant IL-2 and IL-7 (R&D Systems, 402-ML/CF & 407-ML/CF), and 50 µM 2-mercaptoethanol in 5% CO2 at 37°C.

MNK3i cells were generated as described in Saini et al. (manuscript posted to BioRxiv)(30). In brief, MNK3 cells were transduced with pLenti CMV rtTA3 Blast (Addgene plasmid #26429), selected with 10 µg/ml Blasticidin S HCL (Gibco), transduced with a TRE3G-dCas9-KRAB-P2A-mCherry lentivirus, and subcloned (30). MNK3i cells were maintained in MNK3 media plus 10 µg/ml Blasticidin S HCL. MNK3i+sgRNA cells were additionally cultured with 2 µg/ml Puromycin Dihydrochloride (Gibco) beginning 36-48 hr after transduction with sgRNA lentivirus. All cells tested negative for mycoplasma.

### CRISPRi Screen

The mouse genome-wide CRISPRi-v2 library was acquired from Addgene (Addgene #83987-top 5 sgRNAs per gene)(25). This library contains 107,415 sgRNAs that target 20,003 genes and includes 2,170 non-targeting controls. The library (125 µg) was transformed into MegaX DH10B electrocompetent *E. coli* (Invitrogen) and purified from cultures using ZymoPURE II Plasmid MaxiPrep columns (Zymo Research). The amplified library was transfected along with pMD2.G (Addgene #12259) and psPAX2 (Addgene #12260) into HEK293T cells (Mirus TransIT-293T transfection reagent, Mirus) to produce lentivirus. HEK293T supernatant containing the lentivirus was collected two days after transfection and strained with a 0.45 μm filter. The titer of the packaged sgRNA libraries was determined according to published protocols (26).

For library transduction, MNK3i cells were seeded in 15-cm^2^ plates. Three days after seeding (approximately 80% confluent) the pooled sgRNA lentiviruses together with 8 µg/ml polybrene (Millipore Sigma) were added at an MOI of 0.3 and cultured overnight. After 48 hr, the transduced MNK3i cells were selected in 2 µg/ml puromycin for 2.5 days. Puromycin selected cells were treated with 2 µg/ml doxycycline (dox) for 1.5 days, followed by stimulation with 10 ng/ml IL-1b and 10 ng/ml IL-23 (R&D Systems, 401-ML-010/CF & 1887-ML-010/CF) for 21 hr. Stimulated cells received monensin (GolgiStop, BD Life Sciences) during the final 6 hr. The cells were rinsed with 1X DPBS (Gibco), trypsinized (Thermo Fisher), fixed and permeabilized (BD Life Sciences cytofix/cytoperm kit), then stained with 1:100 anti-IL-22 (clone 1H8PWSR, eBioscience) and 1:100 anti-IL-17F (clone eBio18F10, eBioscience) conjugated monoclonal antibodies (eBioscience). Sorting for populations expressing various levels of IL-22 was performed on a FACS Aria III instrument (BD Life Sciences). Cells were double gated for singlets, then on mCherry, and sorted into negative and high bins of IL-22 expression. Of the positive IL-22 cells, the top ∼20% were sorted into the high bin. Post-sort analysis confirmed sorting purities. Another population was not sorted, but rather directly lysed to be a control for library representation.

Genomic DNA was isolated from each population and guide sequences were PCR amplified as published (25, 26). Specific oligonucleotides used for guide generation are listed in Table S1. PCR products (approximately 270 bp) for each population were run on a 1.5% agarose gel, excised, and purified (QIAquick Gel Extraction Kit, Qiagen). Each DNA pool was quantified on a bioanalyzer and sequenced via NGS on a MiSeq (Illumina, Washington University in St. Louis, St. Louis, MO, USA) to assess quantity for correct ratio pooling prior to full-scale NGS analysis on a NovaSeq 6000 (Illumina, Nationwide Children’s Hospital Institute for Genomic Medicine, Columbus, OH, USA). Sequencing of the unsorted population verified even distribution within the guide library.

### Computational analysis of the CRISPRi screen

FASTQ files were trimmed for Illumina Universal Adapters and low-quality bases, then were aligned to the 106,516 unique (of 107,415 possible) 20 bp sgRNA library by the Model-based Analysis of Genome-wide CRISPR/Cas9 Knockout (MAGeCK, version 0.5.9.5) algorithm’s “count” command (31). To focus on sgRNA targeting expressed genes, the raw count file was cross-referenced with a list of genes showing an average transcript per million (TPM) of ≥2.5 in RNAseq data from MNK3i+scramble (sgSCR) cells treated with dox (48 hr) and stimulated with 10 ng/ml IL-1b/23 (21 hr). This curated “expressed” gene library resulted in 48,560 sgRNA targeting 9,262 genes. MAGeCK’s “test” command was applied to generate normalized (method = total) sgRNA and gene-level rankings using Robust Rank Aggregation (RRA). Comparisons featured sgRNAs rescued from sorted IL-22^neg^ versus the IL-22^hi^ populations. Snake plots were generated in RStudio using ggplot2.

### Validation of individual candidates from the CRISPRi screen

Candidate significance was determined by percentile, with the top 2.5% of gene-level (rank) candidates that also had at least one sgRNA in the top 10% by p-value and by log2 fold change (L2FC) being considered for further analysis. Fourteen genes representing a diverse selection of factors (including known regulators) were selected for validation after a literature search. Individual guide sequences for validation of candidate genes were selected from the library based on enrichment in the IL-22^neg^ versus IL-22^hi^ population. The guides of interest and their reverse complements were synthesized (Sigma and IDT), annealed, and ligated into the Esp3I site of sgOpti (Addgene 85681) according to previously published protocols (32). Sanger sequencing of individual plasmids confirmed insert integrity (Ohio State Comprehensive Cancer Center Genomics Core, Columbus, OH, USA). Plasmids containing guides or a SCR control were packaged into lentiviral particles and transduced into MNK3i cells as described above. Two days later, 2 µg/ml puromycin was added to select for lentiviral integrants. Each MNK3i line was treated with 2 μg/ml dox (48 hr), then stimulated with 1 ng/ml IL-1b/IL-23 (21 hr). GolgiStop was added for the final 6 hr. Cells were rinsed with 1X DPBS, trypsinized, and counted by Vi-Cell (Beckman Coulter) to prepare 100K cells for flow cytometry as described above. The percentages of intracellular IL-22 and IL-17F positivity were measured on BD FACSCanto II or BD FACSymphony (BD Life Sciences). Flow cytometry data were analyzed using FlowJo software.

### mRNA analysis

RNA isolation was performed with Trizol reagent (Ambion). Reverse transcription of 250-500 ng RNA was performed using the Verso cDNA synthesis kit (ThermoFisher). qPCR primers are listed in Table S2, and were selected from either Origene or designed by NCBI Primer Blast. For RT-qPCR, 1.5 μl of 7.5X diluted cDNA was used with iTaq universal SYBR supermix (BioRad), 5 μl SYBR, 3 μl nuclease free H_2_0, and 0.25 μl of 10 µM forward & reverse primers. Samples were run in BioRad CFX384, CFX Connect, or C1000 thermocyclers and normalized to beta actin (*ActB*). For nascent transcript experiments, dox-treated cells were incubated with 0.2 mM 5-ethynyl Uridine (EU) for 6 hr followed by RNA isolation using Trizol. De novo RNA expression was measured according to manufacturer’s protocol using the Click-iT™ Nascent RNA Capture Kit (ThermoFisher). For RNA splicing analyses, cells were incubated with 5 nM Pladienolide B (Caymen Chemicals) for 21 hr. cDNA was generated as above. Genomic DNA was used as an unspliced control. PCR was performed with CloneAmp HiFi PCR Premix (Takara Bio) with 5 μl premix, 3 μl H_2_0, 0.25 μl of 10μM forward & reverse primer, and 1.5 μl template. For RNA half-life experiments, cells were incubated with 1 µg/ml Actinomycin D for 0, 6, or 24 hrs. cDNA was generated as above from 500 ng RNA and compared with the untreated sample.

### RNAseq

MNK3i lines were treated with 2 µg/ml dox (48 hr) and stimulated with 10 ng/ml IL-1b/23 (21 hr), resuspended in Trizol, and submitted to GENEWIZ by Azenta Life Sciences (Burlington, MA, USA) for sample extraction, ribodepletion, library preparation, and 2×150 sequencing on a NovaSeq 6000. FASTQ files were concatenated, trimmed for quality and Illumina adapters using Trim Galore, and mapped to GRCm39 using HISAT2. SamTools was used to generate BAM and bigwig files and Rsubread was applied to count the reads. The data were normalized using DESeq2 (negative binomial distribution) or stringtie (TPM). For DESeq2, a cutoff was first applied to remove genes with an average of <15 raw counts per sample. The cutoff for defining differentially expressed genes (DEGs) was an absolute L2FC ≥1.0 and an adjusted p-value ≤0.05. Pathway analyses were performed via DAVID (33, 34) and Enrichr (35, 36, 37). Online platforms NASQAR (38) and VolcaNoseR (39) were used to make principal component analysis and volcano plots, respectively.

### Western blotting

Cells were washed with cold 1X DPBS and lysed using Triton-X lysis buffer with protease inhibition cocktail (cOmplete™, EDTA-free Protease Inhibitor Cocktail, Sigma) and PhosSTOP (Roche). Lysates were agitated for 20 min at 4°C and sonicated on a Biorupter Pico (Diagenode) using the low setting for seven cycles of 30 sec on, 30 sec off. Following addition of 6X Laemmli buffer with 2-mercaptoethanol (ThermoFisher), samples were heated at 95°C (5 min), and 15-50 µg of lysate was loaded into 4-15% precast SDS-PAGE gels (Mini-PROTEAN® TGX™, BioRad). Fractionated proteins were transferred to PVDF membranes (100V, 2 hr, 4°C). Membranes were blocked at ambient temperature in 5% skim milk in 1X TBST (1 hr). Primary antibody was applied overnight at 4°C in 5% skim milk in 1X TBST. Secondary antibody was applied at ambient temperature in 5% skim milk in 1X TBST (1 hr). Blots were incubated in Clarity™ Western ECL Substrate (BioRad) for 5 min and then imaged on a Chemidoc Touch or Chemidoc Go (BioRad). Stripping between antibodies was done using RestorePLUS Western Blot Stripping Buffer (ThermoFischer) for 10-15 min at ambient temperature while being agitated. Stripped membranes were re-blocked for 30 min at RT with 5% skim milk in 1X TBST prior to further staining. Membranes were not stripped prior to a new antibody if the size distance between targets was sufficient and the original antibody demonstrated specificity. Antibodies and concentrations are shown in Table S3.

### Statistical analysis

The authors were not blinded to the samples and no randomization occurred. CRISPRi screen statistics and significance were calculated by MAGeCK as described above. Unless otherwise indicated, data are shown as fold change relative to the sgSCR MNK3i line. Statistics were calculated in GraphPad Prism using one-or two-way analysis of variance (ANOVA) with Dunnett’s multiple comparison test or Student’s t-test (unpaired) as described in the figure legends.

### Data availability

The RNAseq data are deposited to the NCBI Sequence Read Archive (SRA) repository under PRJNA1251803. The CRISPRi screen raw sgRNA counts are provided in Table S4. Materials are available upon reasonable request from the corresponding author.

## RESULTS

### Genome-wide CRISPRi screen for factors controlling type III cytokine expression

To identify novel factors that regulate type III cytokines, we applied a genome-wide CRISPR inhibition (CRISPRi) approach to the murine ILC3-like cell line, MNK3. Primary ILC3 cells are sparse and fastidious ex vivo, but MNK3 cells recapitulate primary ILC3s both epigenetically and transcriptionally, including their expression of type III transcription factors and cytokines (14, 30, 40). To equip the MNK3 line for CRISPRi screening, it was engineered for dox-inducible expression of a dCas9 fused to the potent chromatin repressor, KRAB; heretofore referred to as the MNK3i cell line (Fig. 1A) (30). MNK3i robustly expressed dCas9-KRAB protein upon dox treatment, with minimal, but detectable expression in untreated cells (Fig. 1B). To confirm efficient repression of target genes in MNK3i, we infected cells with two lentiviral expression vectors harboring distinct sgRNAs specific for the *Il22* promoter. Following dox treatment and IL-1b/23 stimulation, the cells containing the sgRNA targeting the *Il22* promoter expressed substantially lower levels of IL-22 protein and *Il22* mRNA compared with MNK3i cells expressing a scrambled guide (Figs. 1C and 1D). In keeping with low level baseline expression of dCas9-KRAB in MNK3i, we also observed a reduction of IL-22 expression in cells containing the *Il22* sgRNA with no dox treatment. As expected, expression of another type III cytokine, IL-17F, was unaffected by the *Il22*-specific sgRNA at either the protein or transcript level (Figs. 1C and 1E).

**Figure 1.**
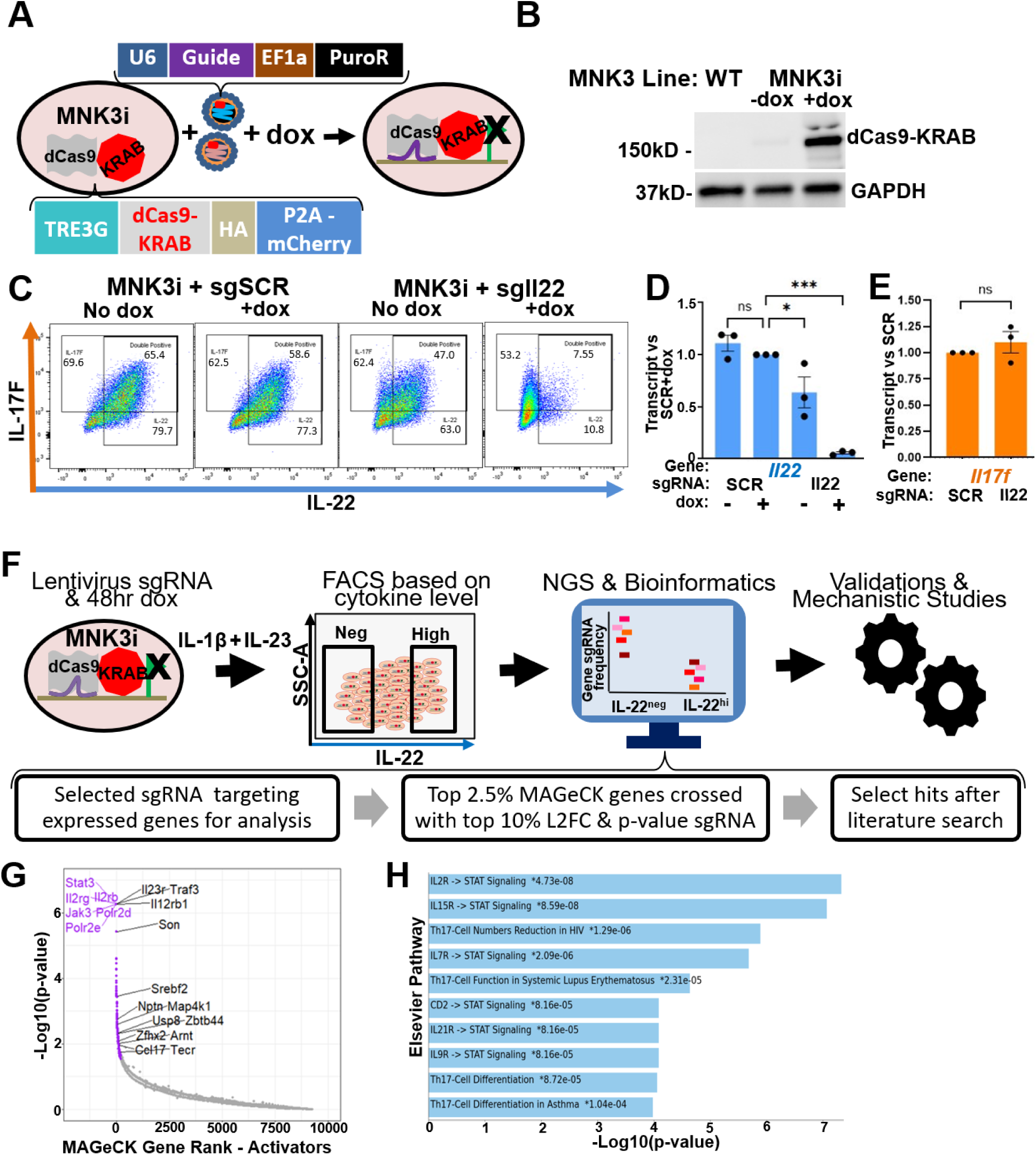
CRISPRi screen identifies known and potential novel regulators of type III cytokines. A) Overview of MNK3i cell engineering. MNK3 cells were stably transduced with a tetracycline (dox) inducible expression vector for catalytically dead Cas9 (dCas9). A MNK3i subclone was then transduced with lentiviral expression vectors for sgRNAs to inhibit specific genes. B) Inducible dCas9 expression in MNK3i cells. Parental MNK3 and MNK3i cells (+/- 48 hr of 2 μg/ml dox treatment) were subject to Western blot for Cas9 and GAPDH (loading control). Representative of n=2. C) MNK3i cells containing either a scrambled guide (sgSCR) or a guide targeting the *Il22* promoter (+/- 48 hr 2 μg/ml dox) were stimulated with 10 ng/ml IL-1b/23 for 21 hr and collected for flow cytometric analysis of IL-17F and IL-22 expression. A representative experiment (n=3) is shown, with numbers indicating total IL-22 % positive, total IL-17F % positive, or % double positive. D-E) MNK3i containing either SCR or *Il22* sgRNA (+/- 48 hr 2 μg/ml dox) were stimulated with 10 ng/ml IL-1b/23 for 21 hr and RNA was collected for RT-qPCR. Data were normalized to *ActB* and shown as fold change relative to the dox-treated sgSCR sample. F) Schematic of the CRISPRi screen workflow. G) CRISPRi results for positive regulators among expressed genes, sorted by gene rank (MAGeCK). Purple dots indicate the top 2.5% of genes. Genes selected for validations are labeled in black. H) DAVID Elsevier pathway analysis of the top 2.5% of genes sorted by gene rank as positive regulators. *p < 0.05, ***p < 0.005, ns = non-significant by one-way ANOVA with Dunnett’s multiple comparison test or unpaired Student’s t-test. Error bars indicate +/- SEM.

An overview of the CRISPRi screen is shown in Fig. 1F. A pre-designed mouse CRISPRi library consisting of 107,415 sgRNAs targeting 20,003 genes (∼5 sgRNAs per target gene) was lentivirally transduced into the MNK3i cell line at an MOI of 0.3 (25). MNK3i cells expressing an sgRNA were selected with puromycin for stable integration of the transduced vectors. The total number of transduced cells was calculated to yield ∼500 stable integrants for each guide in the library. Following dox treatment, the pooled MNK3i cells were stimulated with IL-1b/23, stained for intracellular IL-22/17F, and FACS sorted into IL-22^neg^ or IL-22^hi^ populations (top 20% of positive cells). Genomic DNA from the sorted cell populations was amplified with primers specific to sgRNA-flanking sequences, and the rescued guides were sequenced. Lower capacity sequencing of the sgRNA pool confirmed robust distribution of rescued guides prior to deep sequencing.

Sequencing data for sorted populations were processed using MAGeCK to identify targets depleted or enriched in the respective sorted cell populations (31). The overall quality and distribution of the rescued sgRNA sequences indicated >91% mapped to the original library and there was a low percentage of zero counts (∼0.2%). Importantly, distribution of the rescued guides was well within quality parameters as determined by MAGeCK, with each Gini index for the unsorted, IL-22^hi^, and IL-22^neg^ populations less than 0.08 (desired ≤0.1). MAGeCK analyses were performed at both the individual sgRNA and gene levels (multiple sgRNAs targeting a given gene), and we chose the latter to reduce false positives. We focused additional analyses on genes that were expressed in MNK3i cells at a reasonable level based on RNAseq data (TPM ≥ 2.5). A compendium of results from these analyses is provided in Tables S4 (MAGeCK raw sgRNA counts, unfiltered) and S5 (MAGeCK gene level analysis of expressed targets for IL-22^hi^ versus IL-22^neg^). After MAGeCK normalization (method = total), gene targets enriched in the IL-22^neg^ versus IL-22^hi^ population were considered potential positive regulators of IL-22 expression (Fig. 1G), while those depleted in the IL-22^neg^ versus IL-22^hi^ population were considered potential negative regulators (Fig. S1A).

To highlight the effectiveness and limitations of the screen, we focused on several control genes that were known positive regulators of type III cytokine expression. As expected, sgRNAs targeting promoters for such genes scored highly by MAGeCK, including IL-23 receptor components (IL12RB1/IL23R) and the transcription factor STAT3. Surprisingly, *Il22* itself was not in the top gene results, likely reflecting an overall lower quality of sgRNAs in the library targeting the IL-22 gene compared with the guide we designed specifically for the *Il22* promoter. Indeed, only one of the five library *Il22* sgRNAs was enriched in the IL-22^neg^ population with a log2 fold change (L2FC) ≥1.5, which we validated in subsequent experiments (see below). However, overall, pathway analysis of the top 2.5% of screen hits by gene rank had strong enrichment for Th17-related pathways and STAT signaling (Fig. 1H), which further supported the capacity of our screen to detect factors regulating type III cytokines. As expected in CRISPRi screens, some general cellular factors for housekeeping functions (e.g., splicing and RNA polymerases) were also among the top hits (Fig. S1B).

### Validation of selected candidate genes

To further validate the CRISPRi screen, we selected a set of candidates for focused testing with gene-specific sgRNAs (highlighted in Fig. 1G). In addition to several known factors that control type III cytokines, we selected from new candidates that were: (i) potential positive regulators of IL-22, (ii) in the top 2.5% by gene rank, (iii) contained at least one sgRNA in the top 10% by L2FC and p-value, (iv) not previously linked directly to IL-22 expression, and (v) not general transcription and/or splicing factors (e.g., RNA Pol II subunits). Specifically, we selected sgRNAs for each candidate gene that were enriched in the IL-22^neg^ population by a L2FC of at least 1.5 (Fig. 2A), suggesting their target gene was important for optimal cytokine expression. All genes were expressed at the mRNA level in our MNKi model (Fig. S1C). The impact of each candidate on IL-22 expression was tested by creating a MNK3i line expressing 1-2 guide(s) from the CRISPRi library that targeted the selected gene (Table S1). In brief, individual guide(s) for each selected gene were cloned into a lentiviral vector, transduced into MNK3i cells, and selected for stable integration by puromycin treatment. The selected cells for each set of guides were treated with dox for 48 hr prior to limited stimulation with IL-1b/23 for an additional 21 hr. GolgiStop was added for the last 6 hr. Intracellular IL-22/IL-17F expression was assessed via flow cytometry and RT-qPCR.

**Figure 2.**
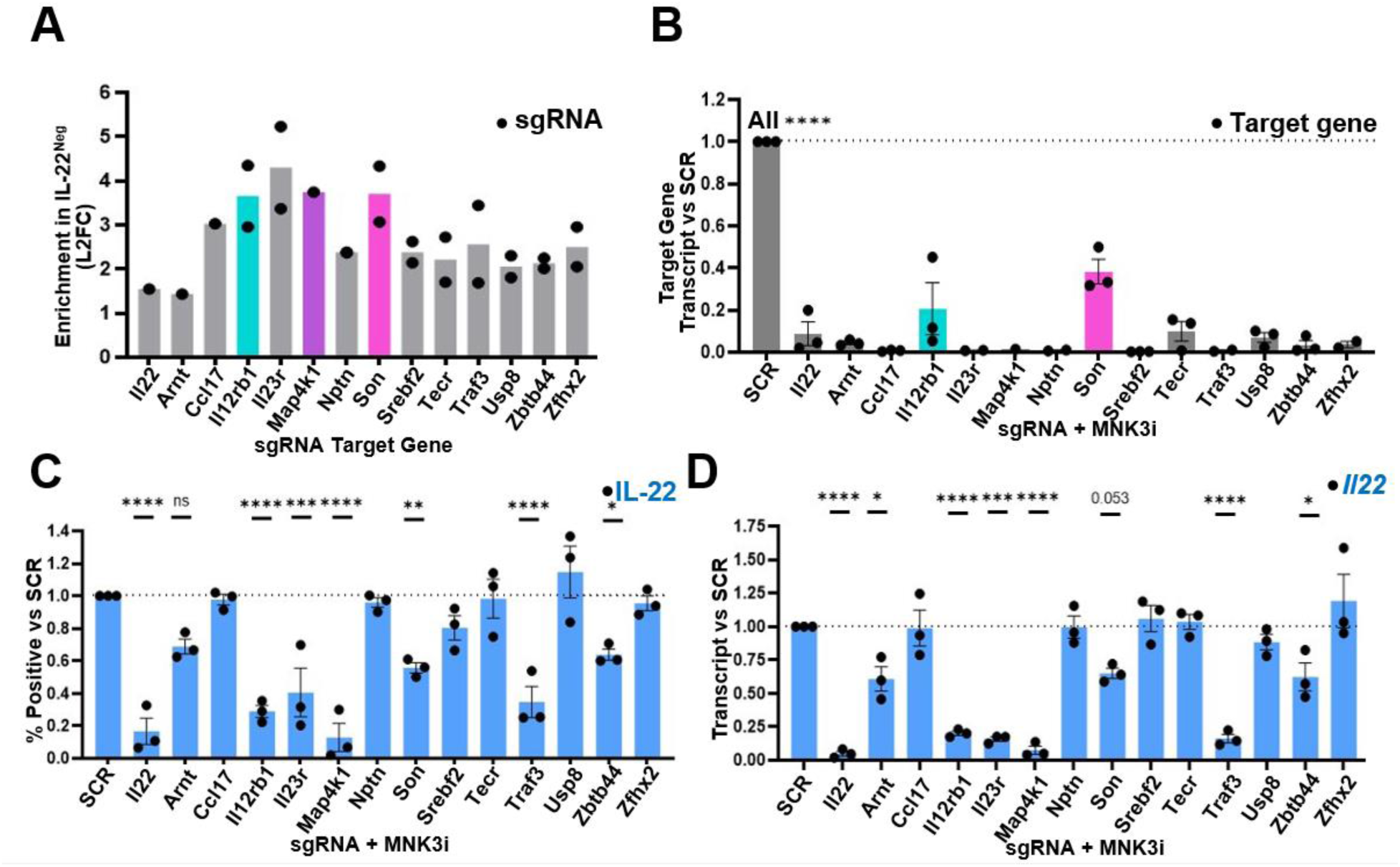
Validation of candidates from the CRISPRi screen. A) Log2 fold change (L2FC) enrichment in the IL-22^neg^ versus IL-22^hi^ populations for sgRNAs corresponding to genes selected for further validation. Follow-up experiments (below) were performed with MNK3i cells containing the indicated sgRNAs for *Il12rb1* (teal), *Map4k1* (purple), and *Son* (magenta). B-D) MNK3i cells were treated with 2 μg/ml dox for 48 hr and stimulated for 21 hr with 1 ng/ml IL-1b/23 + 6 hr GolgiStop. Transcript levels of CRISPRi targeted genes (B) and *Il22* (D) were normalized to *ActB* and shown as fold change relative to sgSCR. Intracellular IL-22 protein (C) was measured by flow cytometry and shown as percent positive relative to sgSCR. *p < 0.05, **p < 0.01, ***p < 0.005, ****p < 0.001, ns = non-significant by one-way ANOVA with Dunnett’s multiple comparison test. Error bars indicate +/- SEM.

As shown in Fig. 2B, all of the lines with specific sgRNAs suppressed their target genes upon dox treatment, with decreases ranging from two-to over a hundred-fold at the mRNA level. As we gated on IL-22 for the screen, we focused our validations on the impact of each target gene knockdown on IL-22 levels; IL-17F expression data are provided in Figs. S1D and S1E. Each transduced MNK3i cell line was henceforth named sg*TargetGene*. As expected, the most enriched *Il22* sgRNA from the CRISPRi library specifically repressed IL-22, but not IL-17F. For the other thirteen candidates, eight reproducibly suppressed expression of one or both cytokines at the mRNA, protein, or both levels. Specifically, the validated hits that served as positive regulators of type III cytokine expression were *Arnt*, *Il12rb1*, *Il23r*, *Map4k1*, *Son*, *Traf3*, *Usp8*, and *Zbtb44* (Figs. 2C and 2D, Figs. S1D and S1E). The other five candidates (*Ccl17*, *Nptn*, *Srebf2, Tecr,* and *Zfhx2*) either led to modest repression in some experiments or no substantial repression of either cytokine, and did not reach statistical significance.

Some of the validated hits are consistent with known functions, especially those encoding essential components of the IL-23 receptor (*Il23r* and *Il12rb1*) (12). Likewise, *Arnt* encodes a binding partner of Ahr (also known as HIF1b), which facilitates its nuclear translocation (41, 42). The latter data suggest that, despite stimulation of MNK3i cells with IL-1b/23, Ahr ligands, naturally present in culture media, are required for optimal expression of both IL-22 and IL-17F. All validated hits are shown in Table 1, together with their general function and possible connections to type III cytokine expression.

**Table 1.** List of validated screen hits. CRISPRi tested hits meeting the statistical cutoff for IL-22 and/or IL-17F protein and/or transcript are listed.

| Gene | Protein | Type III Connection? | General Function |
| --- | --- | --- | --- |
| *Il12rb1* | Interleukin 12 Receptor Subunit Beta 1 (IL12RB1) | Yes, IL-23 receptor | IL-23 and IL-12 receptor component (12) |
| *Il23r* | Interleukin 23 Receptor (IL23R) | Yes, IL-23 receptor | IL-23 receptor component (12) |
| *Arnt* | aryl hydrocarbon receptor nuclear translocator (ARNT), Hypoxia-inducible factor 1-beta (HIF1b) | Yes, via Ahr connection (41) | Ahr Binding Partner for nuclear translocation (41, 42) |
| *Map4k1* | Hematopoietic Progenitor Kinase 1 (HPK1), Mitogen-Activated Protein Kinase Kinase Kinase 1 (MAP4K1) | No | Serine/threonine MAP kinase kinase kinase kinase (43, 44) |
| *Son* | SON DNA Binding Protein (SON) | No | Spicing and transcription co-factor (45, 46) |
| *Traf3* | TNF Receptor Associated Factor 3 (TRAF3) | No | Innate receptor signaling factor, E3 ubiquitin ligase (47, 48) |
| *Usp8* | Ubiquitin Specific Peptidase 8, Ubiquitin Specific Protease 8 (USP8) | Yes, Usp8f/fCd4-Cre CD4 cells show increased IL-17 (49) | Deubiquitinase (49) |
| *Zbtb44* | Zinc Finger And BTB Domain Containing 44 (ZBTB44) | No | Little known. Predicted transcription factor (50, 51) |

Two of the validated genes were unexpected for distinct reasons. *Map4k1* encodes hematopoietic progenitor kinase 1 (HPK1), which has primarily been studied as a negative, rather than a positive, regulator of cytokine expression in the contexts of TCR and BCR signaling (43, 44, 52, 53, 54, 55). Strikingly, knockdown of *Map4k1* abrogated IL-22 expression but had minimal impact on IL-17F levels when compared to sgSCR. The second gene, *Son*, encodes for the nuclear SON DNA and RNA binding protein (SON). SON has selective co-factor roles in splicing and chromatin methylation but has not been linked previously to type III cytokine expression (45, 46, 56, 57, 58). For these reasons, we focused additional analyses on *Son* and *Map4k1*.

### SON regulates IL12R**β**1 expression

Specific knockdown of *Son* mRNA expression by CRISPRi reduced both IL-22 and IL-17F levels upon stimulation of MNK3i by the type III agonists IL-1b/23. SON is a ubiquitously expressed, large nuclear protein with DNA and RNA binding domains (45, 46, 59). Prior studies have indicated that SON functions in nuclear speckles as a docking protein and an RNA splicing co-factor for select splice sites (45, 59). SON also regulates de novo transcription of certain genes via its interactions with menin, a co-factor of the MLL methyltransferase complex (46). SON has multiple isoforms and appears to have general targets, as well as some that are cell type-or agonist-specific (46, 60, 61).

Initially, we confirmed that SON protein was expressed at lower levels in sg*Son* compared with sgSCR (Fig. 3A). Moreover, upon induction by IL-1b/IL-23, sg*Son* cells expressed reduced levels of transcripts for *Son, Il22,* and *Il17f* (Fig. 3B). These reductions were also reflected at the protein levels, as measured by median fluorescence intensity (MFI) of intracellular IL-22 and IL-17F (Fig. 3C). While potent knockdown of *Son* has been shown to reduce cell growth and survival (45, 59), the roughly 50% reduction in sg*Son* permitted unimpaired proliferation and viability during the three-day course of our experiments (Fig. S2A and S2B). Given that our sgRNA targets the *Son* promoter, transcripts for all *Son* isoforms examined were decreased (Fig. S2C).

**Figure 3.**
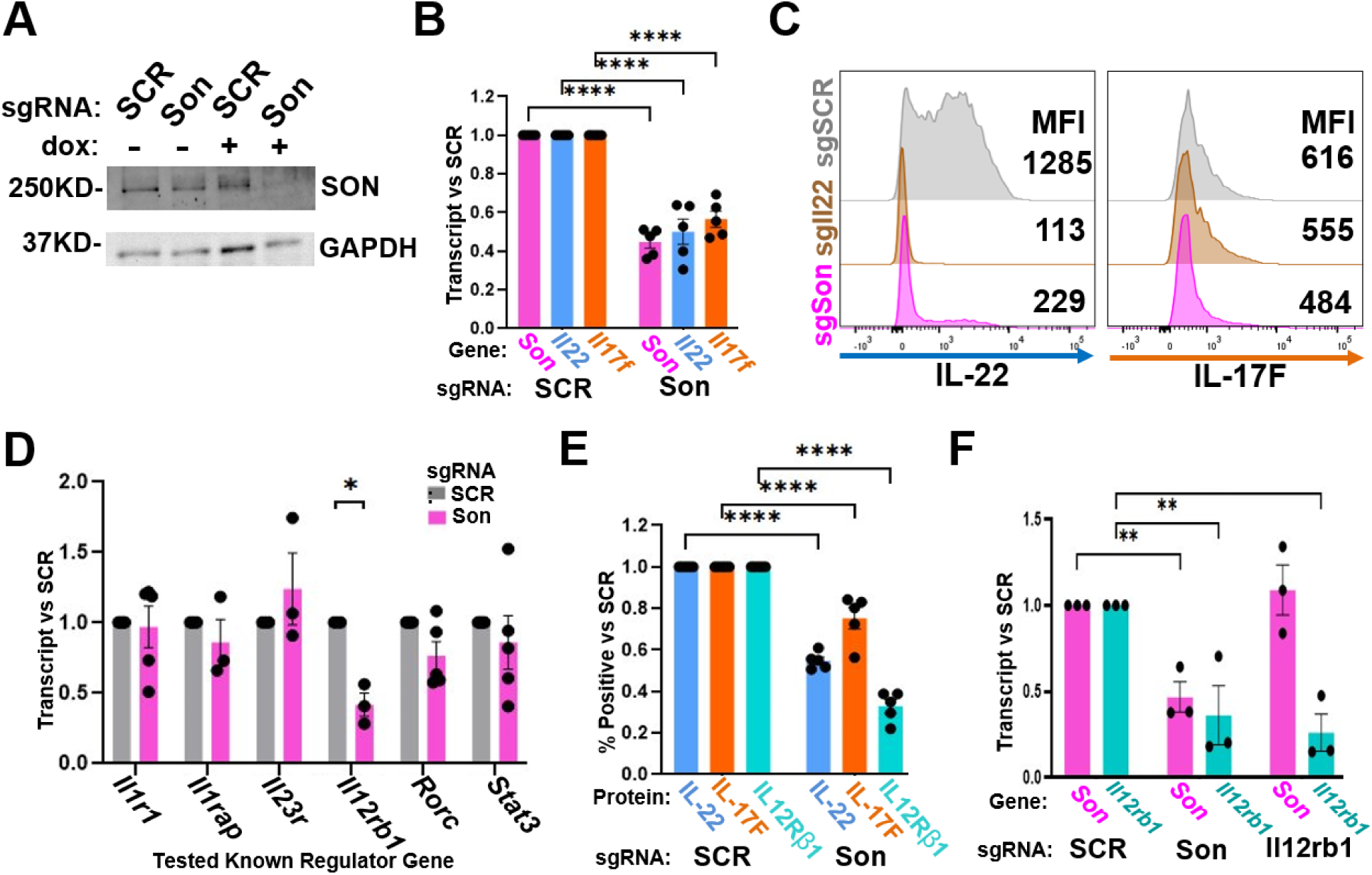
SON promotes optimal IL12Rβ1 expression. A) Whole-cell protein lysates were collected from the indicated MNK3i cell lines (+/- 48 hr 2 μg/ml dox) and analyzed by Western blotting for SON expression. Representative of n=2. B-E) The indicated MNK3i cell lines (+ 48 hr 2 μg/ml dox) were stimulated with 10 ng/ml IL-1b/23 (21 hr + 6 hr GolgiStop). (B) Transcript levels of genes were normalized to *ActB* and shown as fold change relative to sgSCR (n = 5). (C) Flow cytometry median florescence intensity (MFI) for a representative experiment (n = 5). (D) Transcript levels of known IL-22 regulators were normalized to *ActB* and shown as fold change relative to sgSCR. (E) IL12RB1 and intracellular IL-22/17F protein were measured by flow cytometry and shown as percent positive relative to sgSCR. F) Dox-treated (48 hr 2 μg/ml) MNK3i cells were then incubated with 0.2 mM 5-ethynyl Uridine for 6 hr. Nascent RNA was selected for by the Click-iT Nascent Capture Kit. RT-qPCR of nascent transcripts were normalized to *ActB* and shown as relative to sgSCR. *p < 0.05, **p < 0.01, ****p < 0.001, ns = non-significant by two-way ANOVA with Dunnett’s multiple comparison test. Error bars indicate +/- SEM.

To further explore mechanisms by which SON mediates type III cytokine expression, we performed RT-qPCR on a select set of genes involved in pathways upstream of *Il22* and *Il17f*. As shown in Figs. 3D and 3E, SON knockdown resulted in decreased expression of IL12RB1 at the steady state transcript and protein levels, a reduction mirroring the 50% transcript knockdown of SON itself. Notably, the effects of SON reduction are not pleiotropic; transcript levels of other tested receptors (*Il1r1*, *Il1rap*, and *Il23r*) or transcription factors (*Rorc* and *Stat3*) involved in type III cytokine expression were unaffected. Thus, the impact of SON on IL-17/22 expression may be indirect through its regulation of IL12RB1, a core component of the IL-23 receptor.

To determine whether SON depletion affects transcription of the *Il12rb1* gene, we assessed de novo transcripts using the Click-iT assay, in which EU is incorporated into nascent mRNA (62). sgSCR, sg*Son*, and sg*Il12rb1* lines were treated with dox for 48 hr and then incubated with 0.2 mM EU for 6 hr. Nascent transcripts were isolated from total RNA using the Click-iT kit. We found that CRISPRi-mediated repression of *Son* led to a significant decrease in de novo *Il12rb1* and *Son* transcripts when compared with the sgSCR line (Fig. 3F). As a control, repression of the *Il12rb1* promoter specifically reduced its transcripts but not those arising from the *Son* gene. We conclude that SON is required for normal expression of *Il12rb1* at the level of transcription. However, this does not rule out an additional impact on mRNA splicing (see below).

### Transcriptional programs regulated by SON and IL12Rβ1

To further investigate the function of SON in pathways that govern type III cytokine expression, we performed RNA-seq on MNK3i lines during type III stimulation. Specifically, the sgSCR, sg*Son*, and sg*Il12rb1* cells were stimulated (10 ng/ml IL-1b/23) +/- preceding dox treatment. Principal component analysis revealed that dox-treated sg*Son* clustered distinctly compared with sgSCR, while sg*Son* without dox treatment clustered with sgSCR (Fig. 4A). In contrast, even background dCas9 leakage in sg*Il12rb1* (-dox) caused distinct clustering of that line. Hence, we focused our analysis on the dox-treated samples. We first compared transcriptome data from sg*Son* and sgSCR using DESeq2 and found 613 DEGs (L2FC ≥1.0 and adjusted p-value ≤0.05), with 379 upregulated and 254 downregulated upon SON knockdown (Fig. 4B and Table S6). Less than 4.5% of the genes tested were classified as DEGs, which further indicated that SON depletion is not pleiotropically affecting transcriptomes under these conditions. Indeed, the cohort of SON-regulated genes is comparable in number to those affected by IL12RB1 depletion (626 DEGs with 345 up and 281 down, Fig. 4C and Table S7). Notably, sg*Son* and sg*Il12rb1* shared about one-third of their DEGs (206), including *Il12rb1*, *Il22*, *Il17f*, and *Il17a* (Figs. 4D and 4E). The vast majority (93%) of these shared DEGs concordantly changed in both lines compared to sgSCR, with 81 decreasing and 111 increasing (Fig. 4E).

**Figure 4.**
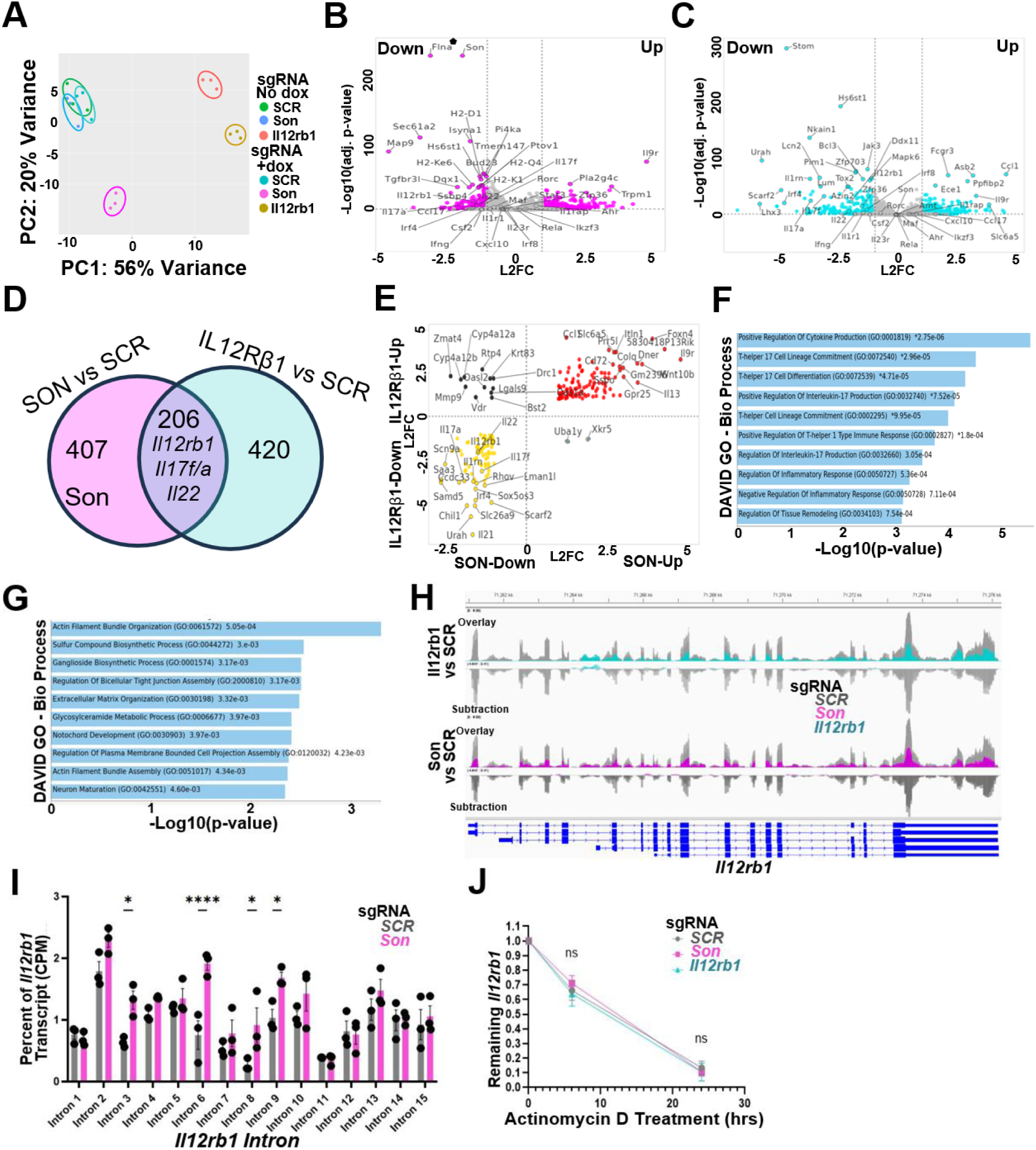
Regulation of genes by SON and IL12Rβ1 under type III activation conditions. A) Principal component analysis of RNAseq data from MNK3i (+/- 48 hr 2 μg/ml dox) stimulated with 10 ng/ml IL-1b/23 (n=3). B-C) Volcano plots of RNAseq for sg*Son* versus sgSCR (B) or sg*Il12rb1* versus sgSCR (C). Differentially expressed genes (DEGs; L2FC ≥ 1.0 and p-value ≤ 0.05) are colored with selected genes labeled. D) Venn diagram showing overlap between sg*Son* and sg*Il12rb1* DEGs. E) Scatter plot of shared sg*Son* and sg*Il12rb1* DEGs plotted as L2FC. F) DAVID Gene Otology - Biological Process analysis of shared directionality sg*Son* and sg*Il12rb1* DEGs. G) DAVID Gene Otology - Biological Process analysis of sg*Son* DEGs not shared with sg*Il12rb1*. H) RNAseq reads mapping to the *Il12rb1* gene visualized in Integrative Genome Browser (IGV). Overlay and subtraction of the sg*Il12rb1* (teal) versus sgSCR reads (grey) and sg*Son* (magenta) versus sgSCR reads (grey) are shown. Subtractions were calculated by bigWigCompare prior to visualization. Samples were group auto-scaled. I) Percentage of Counts Per Million (CPM) normalized total *Il12rb1* reads mapping to each intron in sg*Son* and sgSCR lines. J) *Il12rb1* RNA levels were measured at 6 hr and 24 hr post-treatment with 1 µg/ml Actinomycin D on unstimulated MNK3i cells after initial treatment with 2 µg/ml dox for 48 hr. Results are shown as relative to initial (timepoint 0) *Il12rb1* transcript. ⬟ Indicates DESeq2 adjusted p-value was zero. *p < 0.05, ****p < 0.001, ns = non-significant by two-way ANOVA with Dunnett’s multiple comparison test. Error bars indicate +/- SEM.

The abundance and concordant changes of shared DEGs further supports a primary role for SON in type III cytokine expression through its impact on IL12RB1 expression. The shared DEGs showed enrichment for a large set of type III cytokine-related biological processes, including several linked to Th17 biology and cytokine expression (Fig. 4F). We note that the *Il12rb1* gene is repressed to a stronger degree in sg*Il12rb1* (4.0-fold) than in sg*Son* (2.9-fold). Thus, some of the remaining 420 DEGs specifically identified in sg*Il12rb1* but not in sg*Son* may be rooted in the effectiveness of *Il12rb1* repression. Indeed, sg*Il12rb1*-specific DEGs were primarily linked to cytokine receptor signaling, such as the JAK-STAT pathway (Fig. S2D). By the same logic, the 407 non-shared DEGs for sg*Son* are likely to be independent of IL12RB1 and its downstream signaling pathways. Many genes within this cohort reflect known roles for SON, including its function as a splicing co-factor for genes involved in cytoskeletal organization and neuronal development (Fig. 4G).

As mentioned above, we could not rule out a role for SON in splicing of *Il12rb1* mRNA in addition to its contribution to de novo transcription. Given its function as a splicing co-factor, we leveraged RNAseq data to test this possibility, comparing reads between sgSCR, sg*Son*, and sg*Il12rb1* for retention of *Il12rb1* intronic sequences or differences in alternatively spliced mRNAs. We found that all exons in *Il12rb1* transcripts were reduced in sg*Son* (Fig. 4H) Next, we quantified intronic and exonic reads as a percentage of total *Il12rb1* transcripts to determine whether any of the introns were retained differentially in the sg*Son* line. As shown in Fig. 4I and Fig. S2E, 11 of the 15 introns and all of the exons were present at similar respective levels in sg*Son* versus sgSCR in steady state *Il12rb1* transcripts. However, four of the introns were retained in sg*Son* at significantly higher levels (∼2X). Likewise, SON splicing target *Antkmt* displayed increased retention of introns in sg*Son*, similar to treatment of cells with the RNA splicing inhibitor Pladienolide B, as determined by RNAseq reads and PCR assays spanning a specific exon junction (Figs. S2F and S2G). Thus, even with the ∼50% knockdown efficiency, sg*Son* detectably impacts processing of transcripts sensitive to its function as a splicing co-factor.

Finally, we tested the stability of the *Il12rb1* transcripts, as SON has been shown to help protect select transcripts from nonsense mediated decay (45, 58). However, addition of the transcription inhibitor Actinomycin D (1 µg/ml for 0, 6, or 24 hr) revealed no differences in decay rates for *Il12rb1* transcripts when comparing sgSCR, sg*Son*, and sg*Il12rb1* (Fig. 4J). Hence, these data indicate that splicing of some exon/intron junctions in primary *Il12rb1* transcripts is affected by the SON knockdown, but overall transcript stability is unaffected. Collectively, these experiments demonstrate that SON impacts steady-state levels of *Il12rb1* mRNA, and subsequently protein, through regulation of de novo transcription and RNA processing.

### MAP4K1 is required for IL-22 expression yet dispensable for IL-17

We selected *Map4k1*, encoding the HPK1 protein, as a second hit for downstream experiments due to the powerful impact of its knockdown on IL-22 expression (Fig. 2C and 2D). Conversely, expression of a second type III cytokine, IL-17F, was largely spared (Figs. S1D and S1E). HPK1 is a serine/threonine map 4 kinase predominately expressed in hematopoietic cells (43, 44). This kinase is a negative regulator of the TCR and BCR accessory proteins SLP76 and BLNK, respectively, and has been reported to inhibit dendritic cell activation (52, 53, 54, 55). Accordingly, *Map4k1* knockout mice exhibit increased anti-cancer responses, and T cells deficient in MAP4K1 show enhanced secretion of IL-2, IFNg, and GM-CSF (52, 53). Likewise, NK cells lacking MAP4K1 are marked by increased cytotoxicity, IFNg, and MIP1α (63). Moreover, MAP4K1 was a top validated hit for repressors in CRISPR screens for factors that affect IL-2 or IFNg expression in CD4 and CD8 human T cells (64). In a surprising contrast, *Map4k1* emerged from our screen as a *positive* regulator of IL-22 expression. While only a single *Map4k1* sgRNA showed strong enrichment in the IL-22^neg^ population (Fig. 2A), our validation experiments demonstrated that efficient knockdown of MAP4K1 nearly ablated IL-22 expression (Figs. 2B-2D, Figs. 5A-5C), while IL-17F expression changed only slightly (Figs. 5A-5C, Figs. S1D and S1E).

**Figure 5.**
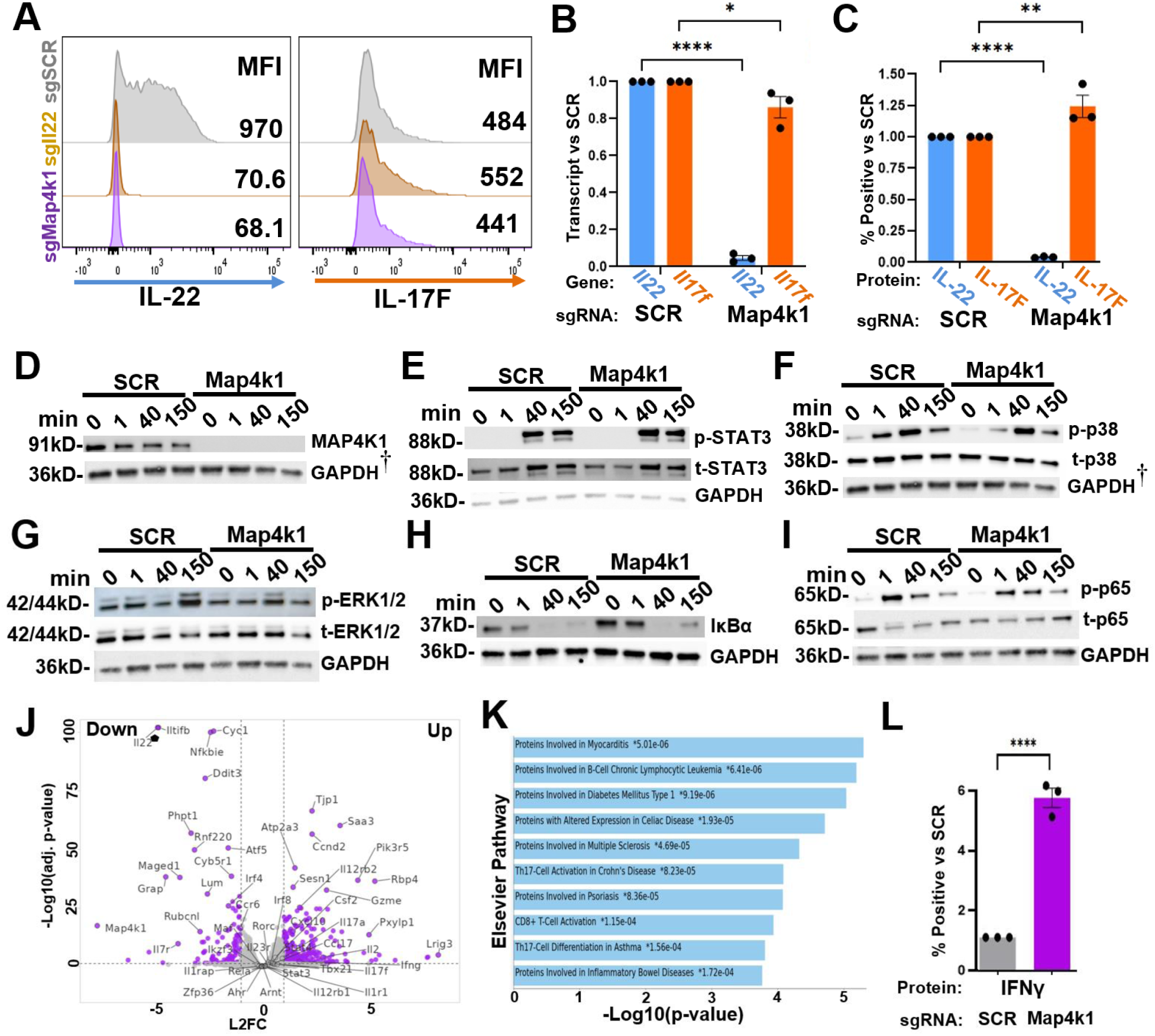
MAP4K1 is required for IL-22 expression with minimal impact on IL-17F. A-C) Dox-treated (48 hr 2 μg/ml) MNK3i cells were stimulated with 10 ng/ml IL-1b/23 (21 hr + 6 hr GolgiStop). (A) Intracellular cytokine MFI for the indicated MNK3i cell lines (representative of n = 3). (B) Transcript levels of cytokines were normalized to *ActB* and shown as fold change relative to sgSCR. (C) Intracellular IL-22/17F protein levels were measured by flow cytometry and shown as percent positive relative to sgSCR. D-I) MNK3i cells were treated with dox (48 hr 2 μg/ml) and stimulated with 10 ng/ml IL-1b/23 for the indicated times. Whole cell protein lysates were analyzed by western blotting with antibodies for the specified proteins. Experiments were performed as biological duplicates, with a representative blot shown for each. Multiple technical replicate blots were run under the same Western conditions. GAPDH was used as a loading control. ^†^ Indicates representative images are from the same technical replicate blot and share the GAPDH (D and F). J) RNAseq Volcano plot for MNK3i cells treated with dox (48 hr 2 μg/ml) and stimulated with 10 ng/ml IL-1b/23 for 21 hr (n = 2). Differentially expressed genes (DEGs; L2FC ≥ 1.0 and p-value ≤ 0.05) are colored with selected genes labeled. K) DAVID Elsevier pathway analysis of sg*Map4k1* versus sgSCR DEGs. L) Dox-treated (48 hr 2 μg/ml) MNK3i cells were stimulated with 10 ng/ml IL-1b/12 (21 hr + 6 hr GolgiStop) and stained for intracellular IFNg. Percentages of IFNg positive cells are reported relative to sgSCR. *p < 0.05, **p < 0.01, ****p < 0.001, ns = non-significant by two-way ANOVA with Dunnett’s multiple comparison test or unpaired Student’s t-test. Error bars indicate +/- SEM.

Prior studies have implicated more distal members of MAPK cascades in IL-23 signaling in ILC3s; however, more receptor-proximal MAPK players in this pathway remain unresolved (65). To examine the impact of MAP4K1 knockdown on select signaling pathways, we stimulated dox-treated sgSCR and sg*Map4k1* cells with 10 ng/ml IL-1b/23 over a 2.5 hr time course and analyzed protein extracts by western blotting. MAP4K1 protein was obliterated in dox-treated sg*Map4k1* cells, including over the entire time course (Fig. 5D). STAT3, the canonical signaling factor downstream of IL23R activation, was robustly phosphorylated regardless of *Map4k1* status (Fig. 5E). Prior studies have shown that inhibition of either p38 (*Mapk14*) or ERK1/2 (*Mapk3*/*Mapk1*) led to significant reductions in IL-22 when splenocytes were induced with IL-23 (65). Although the levels and kinetics of p38 phosphorylation were similar in stimulated sg*Map4k1* cells (Fig. 5F), induced phosphorylation of ERK1/2 was significantly impaired compared with sgSCR controls (Fig. 5G). Thus, although STAT3 and p38 were robustly induced by IL-1b/23 stimulation, neither were able to compensate for the *Map4k1* knockdown.

We also examined the NF-kB pathway, as it is downstream of both IL-1b and IL-23 (66, 67). The kinetics of IκBα degradation and rebound showed no obvious differences between the two lines following IL-1b/23 stimulation (Fig. 5H). Likewise, p65 (RELA) phosphorylation was rapidly induced at similar levels in sgSCR and sg*Map4k1* cells (Fig. 5I). Together, these data indicate that MAP4K1 primarily exerts its effect on IL-22 expression via a pathway involving ERK1/2 phosphorylation.

To elucidate the scope of genes under MAP4K1 control in MNK3, we performed RNAseq of sgSCR and sg*Map4k1* after dox treatment and IL-1b/23 stimulation (Fig. 5J and S2H). Subsequent analysis revealed a DEG cohort of 357 genes (247 up and 110 down; Table S8) with an absolute L2FC of ≥1.0 and an adjusted p-value ≤0.05. In keeping with its known roles as a repressor of immune cell activation, MAP4K1 DEGs were enriched for several chronic autoimmune diseases characterized by hyperactivation of type III and/or type I immune responses, including Celiac Disease, Multiple Sclerosis, Crohn’s Disease, and psoriasis (Figs. 5K and S2I). Likewise, some DEGs impacted by sg*Map4k1* included factors involved in type I and type III polarization and/or effector functions (40). For example, *Il2*, *Ifng*, *Cxcl10*, *Il12rb2* and granzyme transcripts were elevated in sg*Map4k1* cells, while those derived from *Ccr6*, *Irf4*, and *Il7r* were reduced (Fig. 5J).

Of note, we also observed a shift in sg*Map4k1* cells from expressing primarily *Il22* and *Il17f* to co-expression of *Il17* and *Ifng*, a hallmark of T cells that drive pathology in numerous autoimmune diseases (68, 69). Despite this shift, transcript levels of canonical IFNg activating transcription factors, T-bet (*Tbx21*) and *Stat4*, were unchanged in sg*Map4k1* (Fig. 5J). To further confirm that the MAP4K1 knockdown could enhance IFNg protein expression, we stimulated *sgMap4k1* cells with the type I agonists IL-1b and IL-12, given that MNK3 cells exhibit some degree of plasticity (14, 40). As shown in Figs. 5L and S2J, IFNg was increased in *sgMap4k1* versus sgSCR controls after IL-1b/IL-12 treatment. This was consistent with prior work showing MAP4K1 is a negative regulator of IFNg expression (63, 64, 70). Collectively, our findings uncovered an unexpected role for MAP4K1 as a positive regulator of IL-22 but not IL-17 expression, while skewing the type III cells to a pathogenic phenotype co-expressing IL-17 and IFNg.

## DISCUSSION

Cellular pathways that drive the expression of IL-22 and IL-17A/F require a broad array of factors, ranging from cytokine receptors and signal transducers to post-translational modifiers. In the present study, we applied CRISPRi screening to agnostically identify factors required for optimal expression of type III cytokines in an ILC3 model cell line. The screen, which focused on IL-22, successfully identified several known regulators, including STAT3, IL23R, and IL12RB1, as well as novel regulators, such as SON, MAP4K1, and others. While more than half (8/13) of our tested candidates were validated as positive regulators of IL-22 and/or IL-17F expression, the remaining “hits” were false positives (5/13), a level that is consistent with several reported CRISPRi screens, including those with multiple biological replicates (64, 71, 72, 73, 74). Indeed, reproducibility amongst CRISPR screens is an issue noted across the literature, and the number of candidates tested with specific methods for validation is not always provided (75).

Conversely, other candidates may be missed in CRISPRi screens (i.e., false negatives) due to inherent limitations of the experimental systems and/or the sgRNA libraries. Two of these limitations are relevant to the identification of factors controlling type III cytokines. First, CRISPRi targeted genes whose mRNAs or proteins have long half-lives may not be depleted sufficiently to have a significant impact on IL-22 production at the timepoint used for cell sorting. Second, as with all CRISPR-based screens, the quality of the sgRNA library is a major factor. For example, several known genes that should have impacted IL-22 expression did not emerge as “hits” (e.g., *Il22* itself), perhaps due to the low quality of most sgRNAs designed for that gene. In addition, the MNK3 cell model used in our screen derives from NIH-Swiss mice (40), whereas the sgRNA library was designed for the C57BL/6J genome (25). As such, SNPs in the MNK3 genome may have impacted the ability of some guides to target genes efficiently. Notwithstanding these limitations, our CRISPRi screen in MNK3s identified a cohort of genes known to impact type III cytokines, reflecting the overall validity of our approach.

As typical with CRISPR screens, we also identified general factors that regulate mRNA and protein expression, including RNA polymerase components, splicing factors, and those involved in tRNA synthesis. Yet, even broadly expressed genes, like *Son*, can have cell-specific functions that, for example, impact type III cytokine expression. A key finding to emerge from our downstream studies was that SON indirectly regulates IL-22/17 production via being required for optimal expression of IL12RB1, an essential component of the IL-23 receptor. At the mechanistic level, we showed that depletion of SON attenuated de novo transcription of *Il12rb1* and also led to the retention of several introns in steady-state *Il12rb1* transcripts. Together, these data reflect the dual functions of SON as a chromatin-based regulator of transcription and a component of RNA processing machinery that facilitates splicing of certain exons.

Importantly, with regard to *Il12rb1*, the consequences of SON depletion were not pleiotropic, since steady-state transcripts for most other factors involved in the requisite signaling pathways were unaffected. Indeed, RNAseq analysis revealed a large cohort of shared genes that were differentially expressed upon depletion of either SON or IL12RB1, and a second cohort that was specifically affected by SON knockdown. As expected, the shared cohort reflected roles in type III immunity, whereas the latter were mostly related to broader cellular functions. Our discovery that SON functions as a novel regulator of *Il12rb1* expression sets up future studies that will more deeply explore its role in both type I and type III immunity, as well as the underlying mechanisms for SON-mediated control of *Il12rb1* transcripts.

Identification of MAP4K1 as a positive regulator of IL-22 was a surprise, given its known functions as an inhibitor of immune cell activation and cytokine production in multiple cell types. Compared with SON, knockdown of MAP4K1 had a narrower impact on type III cytokines, with a near complete depletion of IL-22 but only a slight reduction of IL-17F. Although the precise mechanisms for this differential effect await further studies, we found that depletion of MAP4K1 primarily impaired stimulation-induced ERK signaling, which is downstream of both type III agonists IL-23 and IL-1b (65, 76). One potential consequence of decreased ERK signaling may be impairment of a given transcription factor that differentially activates *Il22* versus *Il17*. In this regard, Ahr is crucial for ILC3 survival and its deletion in that innate immune subset preferentially attenuated IL-22 rather than IL-17 expression (77). However, during Th17 differentiation, Ahr is required for normal expression of both IL-17 and IL-22 (78, 79). Yet, in MNK3i cells depleted for *Map4k1*, expression of the Ahr target gene *Cyp1b1* was unchanged in our RNAseq data, indicating that Ahr signaling was unimpaired in the sg*Map4k1* line. Collectively, our data further highlights the complexity of regulatory circuits downstream of cytokine receptors that impart each immune cell type with their signature effector functions.

The signaling changes upon MAP4K1 depletion had a surprisingly limited impact on the transcriptome of genes involved in type III immunity. Strikingly, however, some genes involved in type I immunity, including the signature cytokine *Ifng* and the IL-12 receptor component *Il12rb2*, were enhanced upon MAP4K1 knockdown. As a result, the sg*Map4k1* cells co-expressed IL-17 and IFNg following induction with either type III or type I agonists, a hallmark of pathogenic Th17 cells that mediate chronic inflammatory conditions (5). Upregulation of IFNg by IL-17-expressing cells also is observed in the recently described CD26^hi^ T population. These cells, which are often grouped as Th17 cells due to their IL-17 expression, demonstrate more potent anti-cancer capacities than Th1 or Th17 cells (80). Thus, MAP4K1 may regulate yet another aspect of polarization in innate and adaptive lymphoid cells, striking a delicate balance between productive and pathogenic responses. Further elucidation of these pathways may provide new therapeutic avenues for inflammatory diseases, since kinases, like MAP4K1, provide attractive targets for small molecule inhibitors.

Overall, the CRISPRi screen in an ILC3 cell model implicated SON and MAP4K1 as novel regulators of type III immunity through their impact on the expression of signature cytokines (IL-22 +/-IL-17F) and upstream receptors (IL12RB1). Results from our screen provide a rich resource for further studies, including validated positive regulators of IL-22 +/-IL-17F that we have not pursued (e.g., *Traf3*, *Zbtb44*), and negative regulators that need to be further validated. Of special note, the largely unexplored zinc finger protein, ZBTB44, falls into the same category as MAP4K1, in that its depletion impairs expression of IL-22, but not IL-17F, in our ILC3-like model. Future work focusing on factors that govern type III cytokine expression should produce important insights into mechanisms that skew the balance between healthy or damaging immune responses by polarized innate and adaptive lymphocytes.

## Supporting information

https://figshare.com/s/2a6b4f7fafe33ee020fe

## Acknowledgements

We thank Ken Oestreich, Haitao Wen, and Emily Hemann for their gift of Western antibodies and Ning Quan for providing the *Il1r1* qPCR primer sequences. We thank the Genomics Shared Resource Core at The Ohio State University Comprehensive Cancer Center, Columbus, OH for plasmid sequencing.

