## Supplementary material for "CRISPRi Screening Identifies SON and MAP4K1 as Regulators of Type III Cytokine Expression in Innate Lymphoid Cells": https://figshare.com/s/2a6b4f7fafe33ee020fe

**Supplemental Figure 1.** A) CRISPRi screen for negative regulators of IL-22 by MAGeCK gene rank (expressed genes only). Orange dots indicate the top 2.5% of genes. B) DAVID Gene Otology - Biological Process analysis of CRISPRi top 2.5% of positive regulator genes. C) Transcripts per million (TPM) values for selected validation genes (log scale). RNAseq data were derived from sgSCR biological triplicates treated with 2 μg/ml dox for 48 hr and stimulated with 10 ng/ml IL-1β/23 for 21 hr. D-E) IL-17F levels in the indicated MNK3i cell lines treated with 2 μg/ml dox for 48 hr and stimulated for 21 hr with 1 ng/ml IL-1β/23 + 6 hr GolgiStop (n=3). (D) Intracellular IL-17F protein was measured by flow cytometry and shown as percent positive relative to sgSCR. (E) RT-qPCR of *Il17f* normalized to *ActB* and shown as fold change relative to sgSCR. *p < 0.05, **p < 0.01, ***p < 0.005, ns = non-significant by one-way ANOVA with Dunnett's multiple comparison test. Error bars indicate +/- SEM.

**Supplemental Figure 2.** A-B) Viability (A) and cell number (B) of sgSCR and sg*Son* cells as assessed by Vi-Cell after receiving 2 μg/ml dox treatment for 48 hr and stimulation with 10 ng/ml IL-1β/23 for 21 hr (n=7). C) RT-qPCR of *Son* isoforms normalized to *ActB* and shown as fold change relative to sgSCR. D) DAVID Gene Otology - Biological Process analysis of RNAseq differentially expressed genes (DEGs; L2FC ≥ 1.0 and p-value ≤ 0.05) from sg*Il12rb1* versus sgSCR cells treated with 2 μg/ml dox for 48 hr and stimulated with 21 hr 10 ng/ml IL-1β/23 (n=3). E) Percentage of RNAseq counts per million (CPM) mapping to each *Il12rb1* exon. F) RNAseq reads mapping to the *Antkmt* gene in sgSCR (grey) and sg*Son* (magenta) visualized in Integrative Genome Browser (IGV). Biological replicates (n=3) were overlayed and samples were group auto-scaled. G) MNK3i cells received either 5 nM pladienolide B (Plad B) or DMSO (DMSO) for 21 hr or were treated with 2 μg/ml dox (sgSCR, sg*Son*) for 48 hr. RNA was collected for PCR of

*Antkmt* splicing. Genomic DNA (gDNA) was used as an unspliced control. H) Principal component (PC) analysis of RNAseq data from sg*Map4k1* and sgSCR cells treated with 2 μg/ml dox for 48 hr and stimulated with 10 ng/ml IL-1β/23 for 21 hr (n=2). I) DAVID Gene Otology - Biological Process analysis of RNAseq differentially expressed genes (DEGs; L2FC $\geq 1.0$ and p-value $\leq 0.05$) from sg*Map4k1* versus sgSCR cells. J) RT-qPCR of *Ifng* from sg*Map4k1* and sgSCR cells treated with 2 μg/ml dox for 48 hr and stimulated with 10 ng/ml IL-1β/12 (21 hr). Expression was normalized to *ActB* and reported relative to sgSCR. p-value as listed or ***$p < 0.005$, ns = non-significant by two-way ANOVA with Dunnett's multiple comparison test or by unpaired Student's t-test. Error bars indicate +/- SEM.

**Supplemental Table 1.** sgRNA Sequences

**Supplemental Table 2.** RTqPCR and PCR Oligos

**Supplemental Table 3**. Western Antibodies

**Supplemental Table 4**. MAGeCK sgRNA Raw Counts

**Supplemental Table 5.** MAGeCK Gene Level Analysis

**Supplemental Table 6.** RNAseq SONvsScrDESeq2_DEG

**Supplemental Table 7.** RNAseq IL12RB1vsScrDESeq2_DEG

**Supplemental Table 8.** RNAseq MAP4K1vsScrDESeq2_DEG

### Supplemental Figure 1

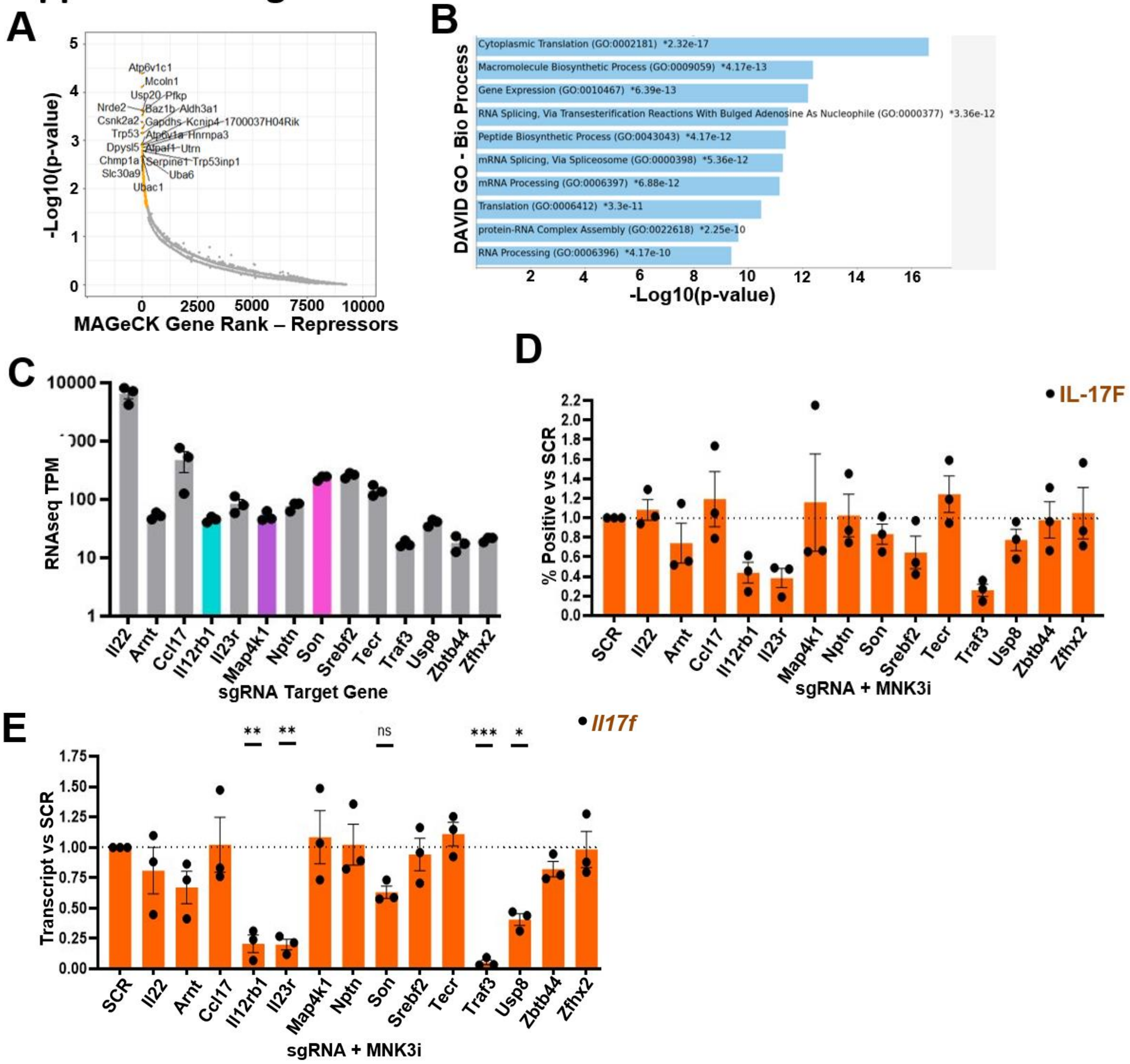

### Supplemental Figure 2

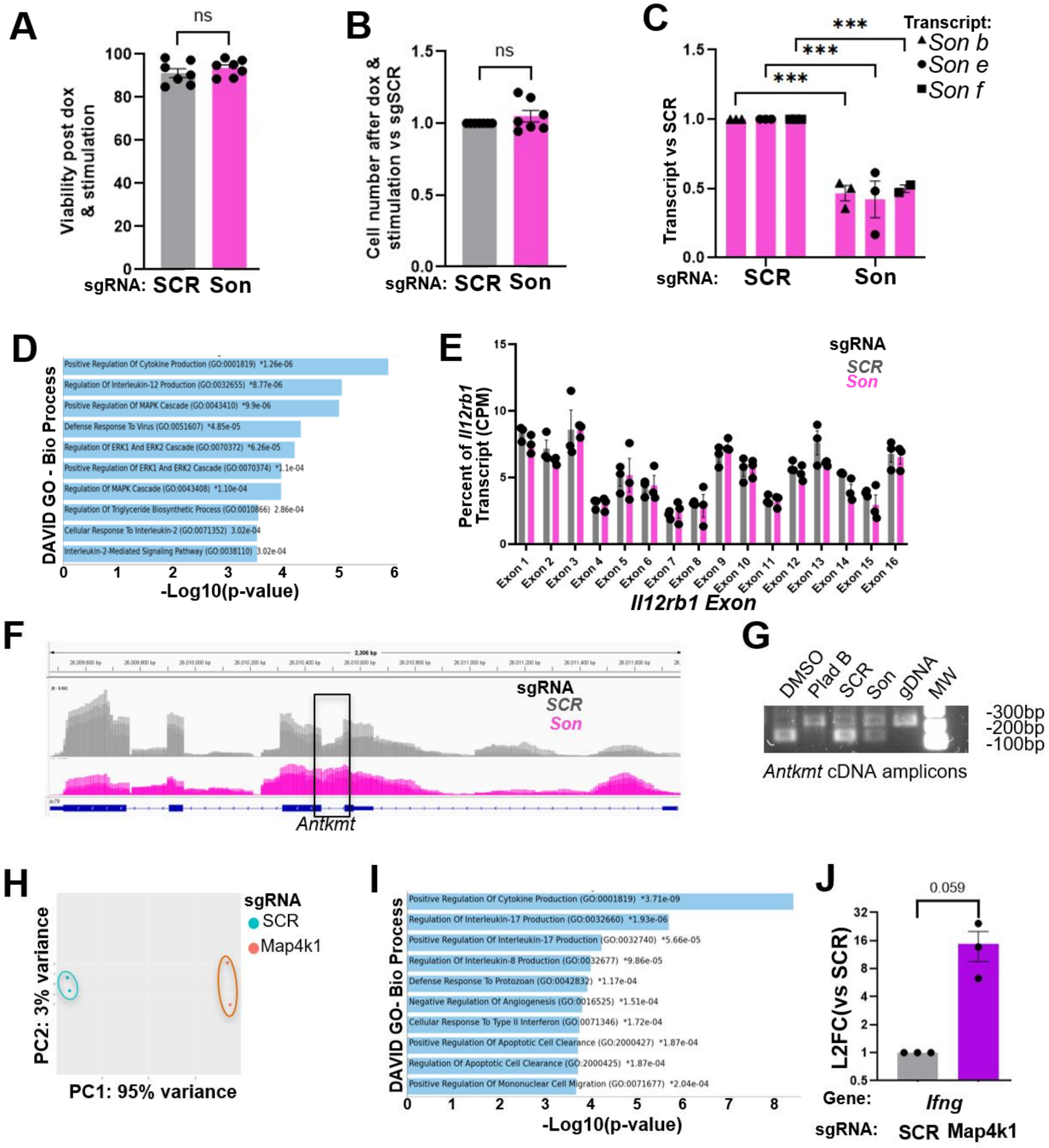

**Supplemental Tables 1 – 8** can be accessed at: https://figshare.com/s/2a6b4f7fafe33ee020fe
